# Thermoresponsive chitosan/silk fibroin/PVA/PVP hydrogel loaded with BDNF promotes functional recovery after stroke

**DOI:** 10.1101/2025.10.20.683259

**Authors:** Mozammel H. Bhuiyan, Emma K. Gowing, Ute Zellhuber-McMillan, Simon F.R. Hinkley, M. Azam Ali, Andrew N. Clarkson

## Abstract

Stroke remains a leading cause of adult disability, driven in part by the formation of a non-permissive extracellular matrix environment that limits endogenous repair. Injectable biomaterials that can modulate this microenvironment while enabling localised therapeutic delivery, represent a promising strategy for post-stroke brain regeneration. Here, we report the development of a thermoresponsive hybrid hydrogel composed of chitosan, β-glycerophosphate, silk fibroin, polyvinyl alcohol and polyvinyl pyrrolidone, engineered to provide a tuneable physicochemical properties and enhanced biological functionalities for intracerebral delivery. Systemic optimisation identified a formulation (F6) that exhibited rapid gelation at physiological temperature, appropriate viscoelastic properties, a microporous architecture, and controlled biodegradation, conducive to cellular infiltration and molecular transport. In a mouse model of photothrombotic stroke, intracerebral delivery of the F6 hydrogel attenuated reactive astrogliosis and microglial activation in the peri-infarct region, while enhancing neurogenesis in the subventricular zone. Notably, incorporation of brain-derived neurotrophic factor within the hydrogel significantly improved functional recovery over 8 weeks, demonstrating the capacity of this system to act as a localised delivery platform for neuro-regenerative therapeutics. Together, this study establishes a tuneable thermoresponsive hydrogel platform that integrates structural support with controlled therapeutic delivery, highlighting its potential as a minimally invasive strategy for modulating the post-stroke microenvironment and promoting functional recovery.

## 1 Introduction

Stroke remains a leading cause of adult disability worldwide, largely due to the limited regenerative capacity of the adult central nervous system [1]. The formation of a non-permissive microenvironment following injury has impeded the development of effective therapies to promote structural and functional recovery after stroke [2]. Current reperfusion therapies including thrombolysis by Tenecteplase and mechanical thrombectomy by self-expanding stent retrievers, can restore blood flow only during the acute phase of stroke [3, 4]. However, due to their narrow therapeutic window, only a small proportion of patients benefit from such treatments, leaving many patients living with permanent disabilities [5–7]. Furthermore, reperfusion therapies do not address the progressive degeneration and impaired repair that occur during subacute and chronic phases after stroke [8]. Consequently, there is a critical need for therapeutic strategies that actively modulate the post-stroke microenvironment to promote lasting tissue regeneration and functional recovery.

A key barrier to regeneration following stroke is the profound remodelling of the extracellular matrix (ECM) which is characterised by the accumulation of inhibitory molecules such as chondroitin sulphate proteoglycans (CSPGs) and the formation of a dense glial scar [2, 9]. These ECM changes suppress axonal growth, limit neural plasticity and restrict endogenous repair processes [10]. Biomaterial-based approaches offer a promising strategy to overcome these limitations by providing structural support while simultaneously modulating biochemical and biomechanical cues within the injured tissue [11, 12]. Among various biomaterials-based strategies, injectable hydrogels have emerged as a particularly promising option for brain repair, as they can be delivered through minimally invasive procedures and readily conform to the irregular geometry of the infarct cavity [13–15]. Following implantation, hydrogels can provide structural support, mimic aspects of the natural ECM and act as a reservoir for localised delivery of therapeutic agents, such as, neurotrophic factors and cells [16–18]. However, despite their promising results in preclinical studies, biopolymer hydrogel-based treatments have yet to be successfully translated into human use. The physicochemical properties of hydrogels, particularly stiffness, porosity, and degradation kinetics, play a critical role in regulating cellular responses, such as inflammation, migration of neural progenitor cells, and tissue remodelling and must be carefully tuned to match the mechanical environment of brain tissue [19, 20]. Therefore, the development of an injectable hydrogel that simultaneously recapitulates the mechanical properties of brain tissue, supports regenerative processes and maintains in situ gelation behaviour, remains a major challenge.

To address these limitations, composite hydrogels incorporating both natural and synthetic polymers have been explored. Thermoresponsive chitosan (CS) / β-glycerophosphate (β-GP) hydrogels are attractive candidates for intracerebral delivery due to their ability to undergo sol-gel transition at physiological temperature, allowing injection as a liquid followed by *in situ* gelation [21–23]. Despite these advantages, CS/β-GP hydrogels alone exhibit limited tunability in mechanical and biological properties, which may restrict their effectiveness in complex neural environments [24]. Incorporation of additional polymers provides an opportunity to modulate the structure of CS/β-GP hydrogels and to overcome these limitations. Previous studies combining other polymers, such as silk fibroin (SF) demonstrated the potential of composite hydrogel in bone tissue engineering by promoting the regeneration of vascularized bone tissue and the deposition of mineralized collagen in a rat model of calvarial bone defects [25]. Similarly, CS/ β-GP/ hyaluronic acid hydrogel was used for the delivery of mesenchymal stem cells into the heart and enhanced cardiac function in a mouse model of myocardial infarction [26]. Silk fibroin (SF), a natural polymer containing cell-adhesive motifs, has been shown to support neural tissue regeneration [27, 28], while synthetic polymers such as polyvinyl alcohol (PVA) and polyvinyl pyrrolidone (PVP) can enhance mechanical stability and viscoelastic behaviour [29–32]. Combining SF, PVA and PVP with CS and β-GP can create a composite or hybrid thermoresponsive hydrogel with both functional motifs and stable mechanical properties for brain tissue repair. However, the combined effects of these polymers blended with thermoresponsive CS/ β-GP hydrogel systems need to be studied to assess their potential for brain tissue repair.

Here, we developed a series of hybrid thermoresponsive hydrogel formulations by blending CS, SF, PVA, PVP and β-GP to tune physicochemical properties that better match the requirements of brain tissue regeneration. We hypothesized that optimising polymer composition would enable the design of an injectable hydrogel capable of modulating the post-stroke microenvironment, attenuating inhibitory glial and inflammatory responses, and supporting endogenous repair processes. We further investigated the potential of the optimised hydrogel to serve as a delivery platform for the key neural plasticity regulator, brain-derived neurotrophic factor (BDNF), to enhance functional recovery after stroke.

## 2 Materials and methods

### 2.1 Materials

CS (Mw: 503 kDa, degree of deacetylation: 91%) was purchased from Shanghai Waseta International Trading Company, China. The molecular weight and degree of deacetylation of CS were characterised using gel permeation chromatography (Agilent PL-GPC 50) and FT-IR spectroscopy, respectively [33]. PVA powder (degree of hydrolysis: 99% hydrolysed, Mw: 89-98 kDa) and PVP powder (Mw: 360 kDa), β-glycerol phosphate disodium salt pentahydrate (Mw: 306.11 Da), lysozyme from chicken egg white, 3-(4,5-dimethylthiazol-2-yl)-2,5-diphenyltetrazolium bromide (MTT), dimethyl sulfoxide (DMSO), and acetic acid were purchased from Merck, New Zealand. Silk fibroin (Mw: 300 kDa) was obtained from Xi’an Herbking Biotechnology Company (Gaoxin District, Xi’an 710075, China) in powder form.

### 2.2 Preparation of hydrogels

#### 2.2.1 Preparation of biomaterials stock solutions

Stock solutions for each biomaterial were prepared as follows: CS 3% w/v, SF 20% w/v, PVA 10% w/v, PVP 10% w/v and β-GP 50% w/v. To prepare CS stock solution, CS was weighed and dissolved in 0.8% (v/v) acetic acid by stirring for 24 h using a magnetic stirrer at room temperature (25°C) to prepare homogenous solutions. To prepare SF and β-GP stock solutions, SF and β-GP were weighed and dissolved in deionized water separately by stirring for 4 h using a magnetic stirrer at 25°C. To prepare PVA and PVP stock solutions, PVA and PVP were weighed and dissolved in deionized water separately by stirring for 4 h using a magnetic stirrer at 80°C.

#### 2.2.2 Preparation of hydrogel formulations

A predetermined volume of stock solution for each biopolymer was mixed in different ratios to prepare hybrid biomaterial solutions as described in **Table 1**. The hybrid biomaterial and β-GP stock solutions were then kept in an ice bath for at least 30 min to cool down to ∼ 4°C. The cold β-GP stock solution was then added dropwise to the hybrid biomaterial solutions to make the final β-GP concentration to 3% w/v with continuous stirring for at least 1 h using a magnetic stirrer to prepare hybrid hydrogel solutions as mentioned in **Table 1**. The final volume of hybrid hydrogel solutions was adjusted by addition of deionized water. When BDNF was added to the hydrogel, the final volume was adjusted so that the concentration of each polymer remains same in the final formulation. The temperature of the system was maintained at ∼ 4°C during the whole process. Finally, the pH was measured using a pH meter (Hanna Instruments pH 209, USA).

**Table 1:** Composition and characteristics of the hybrid hydrogels.

| Name of hydrogels | Concentration (% w/v) | | | | | pH | Gelation time at 37°C (min) | Gelation temp. (°C) | Porosity (μm) Mean ± SEM |
| --- | --- | --- | --- | --- | --- | --- | --- | --- | --- |
| | CS | SF | PVA | PVP | β-GP | | | | |
| F1 | 0.75 | - | - | - | 3 | 7.14 | 5 | 36 | 56.3 ± 1.3 |
| F2 | 0.75 | 3 | - | - | 3 | 7.10 | 26 | 39 | 79.9 ± 4.3 |
| F3 | 0.75 | - | 2 | - | 3 | 7.07 | 1 | 31 | 35.6 ± 1.1 |
| F4 | 0.75 | - | - | 1 | 3 | 7.05 | 3 | 34 | 23.0 ± 17.1 |
| F5 | 0.75 | 3 | 2 | 1 | 3 | 7.05 | 8 | 39 | 18.8 ± 19.5 |
| F6 | 0.75 | 2 | 1 | 0.5 | 3 | 7.04 | 2 | 36 | 30.2 ± 16.6 |

CS: chitosan, SF: silk fibroin, PVA: polyvinyl alcohol, PVP: polyvinyl pyrrolidone, β-GP: sodium β-glycerophosphate.

### 2.3 Characterisation of hydrogels

#### 2.3.1 Gelation time

Gelation time of the hybrid hydrogels was determined by the inverted tube test as described previously [34]. Briefly, the hydrogel forming solution for each hydrogel was added to a transparent glass vial and incubated in a 37 °C water bath. The viscosity of the solution was monitored every 30 s by inverting the tubes horizontally. The time at which the solution stopped flowing was recorded as the gelation time.

#### 2.3.2 Rheological properties

Rheological properties of the hybrid hydrogels were investigated using an oscillatory rheometer (Discovery HR-3, TA Instruments) equipped with a cone and plate geometry of 2° and 60 mm in diameter. Freshly prepared hydrogel-forming solutions were used for the rheological analysis. An oscillatory stress sweep test was performed from 0.1 to 10 Pa with a frequency of 0.1, 1, 10, and 50 Hz at 37 °C to confirm that the frequency and strain were within the linear viscoelastic region (LVR). A frequency sweep test was performed for each hydrogel from 0.1 to 100 rad/s at a constant stress of 1 Pa and at 37 °C. Based on these test results, a stress of 1 Pa and a frequency of 1 Hz was chosen for temperature sweep and time sweep tests. A temperature sweep test from 25 to 50 °C at a heating rate of 1 °C/min and a time sweep test at 37 °C for 60 min were performed for each hydrogel with a constant frequency of 1 Hz and stress of 1 Pa to determine the effect of temperature and time, respectively, on the storage modulus (G’) and loss modulus (G”) of hybrid hydrogels. The G’ represents the elastic property and describes the solid-state behaviour, whereas G’’ characterises the viscous property as seen with the liquid-state behaviour of the sample. The temperature and time at which the G’ value crossed the G” value is considered gelation temperature and gelation time, respectively.

#### 2.3.3 Morphology

The surface morphology along with porosity of the hybrid hydrogels were observed by scanning electron microscopy (SEM) (model: TM3030, Hitachi) as described previously [35]. The freeze-dried hydrogel samples were coated with palladium-gold using an Emitech K575X Peltier-cooled high-resolution sputter coater (EM Technologies Ltd., Kent, England) prior to analysis. The coated samples were viewed at an accelerating voltage of 15 kV. The pore diameter was assessed by measuring Feret’s diameter of 6 pores in three randomly selected sections in each SEM micrograph [29].

#### 2.3.4 Enzymatic biodegradation

The enzymatic biodegradation was measured Gravimetrically as described previously [36]. Briefly, a small cube-shaped sample of freeze-dried hydrogel was cut (1 cm^3^), weighed (W_i_), and immersed in PBS (pH 7.4) containing lysozyme (2 µg/ml) at 37 °C for 1, 3, 5, 7, 14, and 21 days. At each timepoint, the samples were removed from the enzymatic media, freeze-dried and weighed (W_t_). Enzymatic degradation was assessed using the following formula, degradation (%) = (W_i_ – W_t_)/W_i_ X 100.

#### 2.3.5 Cytotoxicity testing

Cytotoxicity was assessed using the MTT assay on PC12 cells (PC12, catalogue no. CRL-1721, ATCC). Briefly, PC12 cells were seeded in 96-well plates at a density of 1 x 10^4^ cells per well in RPMI media (Gibco) supplemented with 10% foetal bovine serum (FBS, Gibco) and 1% penicillin-streptomycin. Following cell adhesion, either of PBS (control), β-GP, biopolymer and hybrid hydrogel solutions were added to the wells in triplicate (**Table 1**). The cells were then incubated for 24 h in an incubator (Heracell VIOS 160i, Thermo Scientific) at 37 °C and 5% CO_2_. Subsequently, media containing testing solutions was replaced by media containing MTT (0.4 mg/ml) and incubated for another 4 h. Following incubation, MTT containing media was removed and formazan was dissolved with DMSO. Finally, absorbance was measured at 550 nm using a microplate reader (SpectraMax i3x MultiMode Detection Platform, Molecular Devices, USA). The average absorbance of control PBS-exposed cells was considered as 100% cell viability with average change in absorbance of β-GP solution biopolymer and hybrid hydrogel treated wells measured to calculate cell viability (%).

### 2.4 In vivo study

#### 2.4.1 Animals

All animal procedures were carried out in accordance with the institutional guidelines and approved by the University of Otago Animal Ethics Committee (AUP# AUP-19-170). Only male mice were used in the studies to minimise variability between experimental groups [37]. Male C57BL/6 mice, aged between 2-3 months, body weights between 25-30 g were purchased from the Hercus-Taieri Research Unit of the University of Otago, New Zealand and acclimatised with their living condition for at least 7 days before surgery. Animals were housed under a 12h light/dark cycle with ad libitum access to food and water. The sample size for each experimental group was determined by a power analysis based on previous similar studies (Table S1). Animals were assigned randomly to experimental groups, and group information was concealed as per recommendations from the Stroke Therapy Academic Industry Roundtable (STAIR), Stroke Recovery and Rehabilitation Roundtable (SRRR) and Animal Research: Reporting of In Vivo Experiments (ARRIVE) guidelines for all behavioural and histological assessments [38–40].

#### 2.4.2 Photothrombotic stroke surgery

Focal cerebral infarction was induced using the photothrombotic method as described previously [41]. In brief, animals were anesthetised using isoflurane (2 – 2.5% in O_2_) and placed in a stereotaxic apparatus. The skull was then exposed through a midline incision and underlying connective tissue, and periosteum cleared and dried. A cold light source (3300KW, Scott, LCD) attached to a 20x objective, giving a 2 mm diameter illumination, was positioned 1.5 mm left from bregma and 0.2 ml Rose Bengal solution (Sigma Aldrich, 10 mg/ml in normal saline) was administered intraperitoneally. After 5 min, the brain was illuminated through the intact skull for 15 min, creating an infarction in the primary motor cortex [41]. The body temperature was maintained at 36.9 ± 0.2°C using a rectal probe attached to a homeothermic blanket during the whole procedure (Harvard Apparatus, Massachusetts, USA).

#### 2.4.3 Intracerebral implantation of hydrogel

Hydrogels were intracerebrally injected into the infarct cavity 5 days after stroke. In brief, mice were anesthetised using isoflurane (2 – 2.5% in O_2_), placed in a stereotaxic apparatus, the skull exposed through a midline incision and cleaned as described before. A small bur hole was created in the skull over the stroke region and 8µl of hydrogel (either of F1, F2, F3, F6) or hydrogel (F6) + BDNF solution was injected into the infarct cavity using a 30-gauge needle attached to a Hamilton syringe (Hamilton®, USA) connected to a pump with an infusion rate of 1 µl/min. The needle was withdrawn 1 min after the injection and the skin glued back together using surgical glue.

#### 2.4.4 Behavioural assessment

The effect of hybrid hydrogel implantation with or without growth factors on behavioural function was assessed using the grid-walking test and the cylinder test as described previously [41]. Behavioural testing was done 1 week before (baseline) and 1, 2, 4, 6 and 8 weeks after stroke surgery and at the same time of each day. For the grid-walking test, each mouse was allowed to walk freely on a 12mm^2^ wire mesh with a grid area of 32 cm X 20 cm for 5 min. Video recordings were analysed offline by an observer blinded to the treatment group, and the total number of normal steps taken, along with the total number of foot faults for each limb counted. For each animal, an average of twenty contacts with the grids were considered for analysis. A foot fault is defined when the animal paw slipped through the grid hole due to inadequate support, or the animal is resting on the grid by placing its wrists on the grid. Percent foot faults were then calculated using the following formula: number of foot-faults / (number of foot-faults + number of non-foot-fault steps) X 100. For the cylinder test, each animal was placed inside a clear plexiglass cylinder apparatus with a diameter of 10 cm and a height of 15 cm and allowed to explore for 5 min. A mirror was placed on one side of the cylinder while a video recorder was positioned on the other side of the cylinder to get a 360° view of the movements while recording. Recorded videos were analysed by playing in slow motion (1/2 of real time speed, QuickTime Player, version 7.7.9) to calculate the total time mice spent on each forelimb during each rear. For each animal, an average of twenty rears were considered for analysis. Percent use of contralateral forelimb was then calculated using the formula given below: average time of right forelimb use / (average time of right forelimb use + average time of left forelimb use + average time of both forelimb use) X 100.

#### 2.4.5 Brain tissue processing

Mice were anesthetised using sodium pentobarbital (7.5 mg/kg body weight, intraperitoneal) and intracardially perfused with 4% paraformaldehyde solution (PFA). The brains were immediately collected and postfixed in 4% PFA over night before being transferred to 30 % sucrose solution in PBS. Brains were cut into 30 µm thick coronal sections using a freezing stage sledge microtome (Leitz Wetzlar, Germany) and the sections were stored in cryoprotectant at – 20°C.

#### 2.4.6 Infarct volume

Infarct volume was determined by histological assessment using Cresyl violet staining, as described previously [42, 43]. For each animal, every sixth section throughout the coronal plane was stained with Cresyl violet and digitised using Stereo Investigator software (Microbrightfield). The edges of the core of the infarct were defined using the drawing tool of Fiji ImageJ (National Institute of Health, USA) and the area of the selected regions were measured by an observer blinded to the animal’s treatment groups. Infarct volume was then quantified as follows: Infarct volume (mm^3^) = Area (mm^2^) x Section thickness (mm) x Section interval.

#### 2.4.7 Immunohistochemistry

Brain sections were washed three times with tris-buffered saline (TBS) at room temperature. Sections were then blocked with 5% normal goat serum (NGS) or normal donkey serum (NDS) in 0.3% w/v Triton X-100 in TBS solution for 60 min with continuous shaking (60 rpm) at room temperature. The sections were then incubated in polyclonal chicken anti-glial fibrillary acidic protein (GFAP) antibody (1:2000, cat. AB5541, Sigma Aldrich) or polyclonal rabbit anti-ionised calcium binding adaptor molecule 1 (Iba1) antibody (1:800, cat. 019-19741, Fujifilm, Japan) or polyclonal goat anti-doublecortin (Dcx) antibody (dilution 1:400, cat. sc-8066, Santa Cruz) diluted in 2% NGS/NDS and 0.3% w/v Triton X-100 containing TBS solution at 4°C for 48 h on an orbital shaker. Sections were then washed in TBS before being transferred to goat anti-chicken Dylight® 488 antibody (1:400, cat. ab96947, Abcam) or donkey anti-rabbit Dylight® 550 antibody (1:800, cat. ab96892, Abcam) or donkey anti-goat Dylight® 549 antibody (1:800, cat. 705-505-147, Jackson ImmunoResearch) diluted in 2% NGS/NDS and 0.3% w/v Triton X-100 in TBS solution for 2 h at room temperature. Following subsequent washes in TBS, sections were incubated in Hoechst solution for 10 min. Sections were then washed in TBS and mounted on gelatine-coated glass slides, air-dried, passed sequentially through alcohol (50%, 70%, 95%, 100%) and xylene, and finally cover slipped using DPX mounting solution.

#### 2.4.8 Image analysis

Photomicrographs of the sections were taken using an inverted microscope (Eclipse Ti2, Nikon, Japan). To quantify the expression of GFAP, the integrated density value (IDV) was quantified across four regions of interests (ROIs, 200 µm x 200 µm): (1) cortical layer 2/3 peri-infarct, (2) cortical layer 5 peri-infarct, (3) cortical layer 2/3 lateral (approximately 600 µm apart from peri-infarct regions), (4) cortical layer 5 lateral using Fiji ImageJ software. To quantify the expression of Iba1, IDV was quantified across two ROIs (200 µm x 200 µm): (1) cortical layer 2/3 peri-infarct, (2) cortical layer 5 peri-infarct. To quantify the expression of Dcx, the raw integrated density value (RIDV) of the subventricular zone, wall of the lateral ventricle was quantified using Fiji ImageJ software.

### 2.5 Statistical Analyses

Statistical analyses were done using GraphPad Prism 9 (GraphPad software Inc., San Diego, USA). Differences between the porosity of hybrid hydrogels were evaluated by one-way analysis of variance (ANOVA). The difference between enzymatic biodegradation of hybrid hydrogels was analysed by two-way ANOVA. Tukey’s multiple comparison test was used to further investigate any significant effect observed. Quantification of infarct volume and immunohistochemical analyses were also assessed by one-way ANOVA and Tukey’s multiple comparison test was conducted to further investigate any significant effect observed. A confidence interval of P < 0.05 was considered statistically significant. All bar graphs are presented as mean ± standard deviation (SD) unless otherwise stated.

## 3 Results

### 3.1 Characterisation of hybrid hydrogel formulations

#### 3.1.1 Hydrogel design

Hydrogels intended for brain repair must be injectable, exhibit rapid gelation under physiological conditions, and possess a porous structure to support cellular infiltration and molecular transport [17]. To achieve these properties, we engineered hybrid hydrogels that remain in a liquid state at room temperature (25_) and undergo rapid sol-gel transition at physiological temperature (37_), enabling minimally invasive delivery and retention within the infarct cavity.

In our previous study, we reported that hydrogels composed of 0.75% (w/v) CS and 3% (w/v) β-GP are porous, non-cytotoxic, and have suitable thermoresponsive gelation properties for injectable applications [44]. Building on this, we combined 0.75% CS and 3% β-GP with SF, PVA and PVP at concentrations determined to be non-cytotoxic (**Fig. S1**), generating six hybrid hydrogel formulations (F1-F6, **Table 1**). These formulations enabled systemic evaluation of how polymer composition influences hydrogel properties relevant to brain tissue engineering.

#### 3.1.2 Viscoelastic and thermoresponsive properties

The viscoelastic behaviour and mechanical properties of the hybrid hydrogels were characterised using a dynamic frequency sweep test [45]. Frequency sweep tests were conducted after 1 h of incubation at 37°C to allow sufficient time for gelation. All hybrid hydrogels showed viscoelastic behaviour characteristic of soft biomaterials, with storage modulus (G′) higher than loss modulus (G”) at low frequencies, indicating dominant elastic behaviour after gelation. Notably, incorporation of 2% PVA (F3) increased G′ relative to the base formulation (F1), suggesting enhanced mechanical stability. In contrast, addition of 3% SF (F2) or 1% PVP (F4) decreased G′, indicating a softer network structure. Addition of SF, PVA, PVP in a single formulation (F5) further reduced G′ compared to the base formulation (F1). The optimised formulation (F6) in which a lower concentration of SF, PVA and PVP were added than formulation F5, showed an increased G′, suggesting a tuning effect of these biomaterials on mechanical stability. At higher frequencies, all formulations exhibited G′ decreased sharply while G” remained relatively unchanged, suggesting oscillatory shear thinning behaviour. This shear-thinning property can be advantageous for injectable applications.

Temperature sweep analysis demonstrated a sharp transition from liquid-like to solid-like behaviour near physiological temperature. The temperature at which G′ surpasses G” is defined as gelation temperature as this marks the transition from liquid to solid state [46, 47]. Gelation temperatures for each formulation are summarised in **Table 1**. Gelation temperature of the hybrid hydrogels ranged from 31°C to 39°C depending on composition. Specifically, SF increased gelation temperature, whereas PVA and PVP lowered it, indicating differential modulation of intermolecular interactions within the hydrogel network. The optimised formulation (F6) exhibited gelation at ∼36°C, ideally aligned to physiological conditions.

#### 3.1.3 Gelation time

Gelation times varied substantially across formulations, ranging from 1 to 26 minutes at 37°C (**Table 1)**. The base formulation (F1) gelled within 5 minutes, while incorporation of SF significantly delayed gelation (F2: 26 min). In contrast, the PVA containing formulation (F3) and the PVP containing formulation (F4) showed rapid gelation (∼1 and ∼3 min, respectively), indicating accelerated network formation. Importantly, the optimised formulation (F6) achieved rapid gelation (∼2 min), balancing injectability with efficient in situ stabilisation. These findings were further validated using a rheological time sweep test, as shown in **Fig. 1A**.

**Figure 1:**
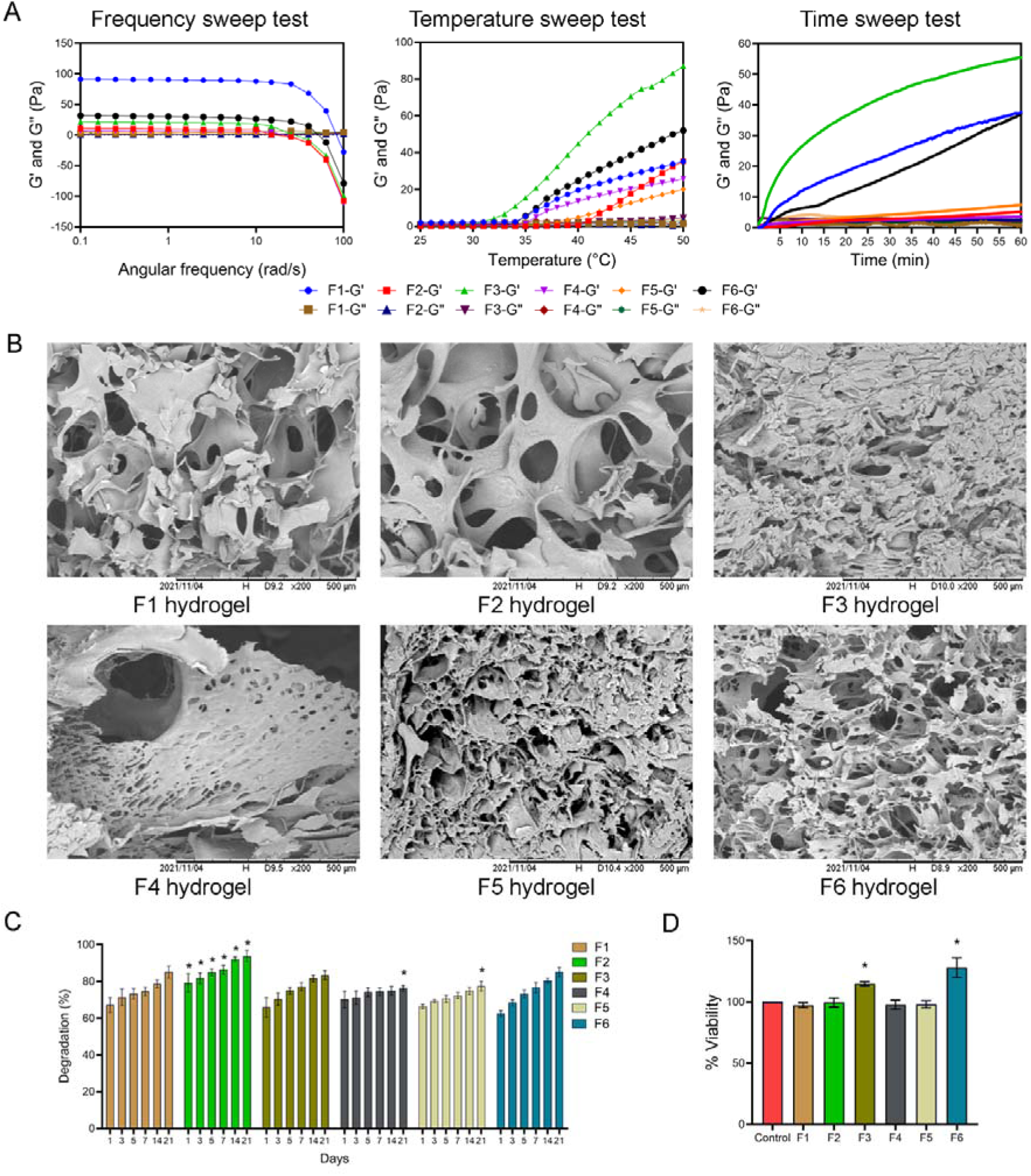
Physical and biological characterisation of hybrid hydrogel formulations. A) Frequency sweep, temperature sweep and time sweep tests of hybrid hydrogels. B) SEM image of the surface morphology of hybrid hydrogels. C) Enzymatic degradation of hybrid hydrogels. Data are expressed as mean ± SEM for n = 3. Data were analysed by two-way ANOVA followed by Tukey’s test for post-hoc comparisons. Asterisks (*) represented significant values. D) Cell viability of PC12 cells grown inside hybrid hydrogels after 24 h. Data are expressed as mean ± SD for n = 3. Data were analysed by one-way ANOVA followed by Tukey’s test for post-hoc comparisons. Asterisks (*) represented significant values. F1: 0.75% chitosan / 3% β-GP, F2: 0.75% chitosan / 3% β-GP/ 3% SF, F3: 0.75% chitosan / 3% β-GP/ 2% PVA, F4: 0.75% chitosan / 3% β-GP/ 1% PVP, F5: 0.75% chitosan / 3% β-GP/ 3% SF / 2% PVA / 1% PVP, F6: 0.75% chitosan / 3% β-GP/ 2% SF / 1% PVA / 0.5% PVP.

#### 3.1.4 Morphology

The interior morphology and porosity of the hybrid hydrogels were examined using SEM. Analysis of SEM images revealed all hybrid hydrogels possessed an interconnected porous structure, with pore size strongly dependent on polymer composition (**Fig. 1B)**. SF-containing hydrogel (F2) exhibited larger pores, whereas PVA– and PVP-containing formulations (F3–F5) displayed more compact structures with reduced pore size. The optimised formulation (F6) exhibited intermediate pore size, suggesting a balance between structural integrity and porosity conducive to cellular infiltration.

#### 3.1.5 Enzymatic biodegradation

Enzymatic biodegradation of the hybrid hydrogels was assessed in vitro using PBS containing lysozyme at 37°C for 1, 3, 5, 7, 14, and 21 days. All formulations demonstrated progressive, time-dependent degradation over time (**Fig. 1C)**. SF-containing hydrogels (F2) exhibited significantly accelerated degradation compared to F1 at early time points, whereas PVP-containing formulations showed modestly reduced degradation at later time points. The F6 formulation exhibited a degradation profile comparable to the base hydrogel (F1), indicating stable yet biodegradable behaviour suitable for in vivo applications.

#### 3.1.6 Cytotoxicity

Cytotoxicity of the hybrid hydrogels was evaluated by MTT assay using PC12 cells. All hybrid hydrogels maintained high cell viability (>90%) following 24 h exposure to PC12 cells, confirming cytocompatibility (**Fig. 1D)**. Notably, F3 and F6 significantly increased cell viability relative to control, suggesting that specific polymer combinations may enhance cellular responses. These results indicate that incorporation of synthetic polymers does not compromise, and may even improve, biological compatibility.

### 3.2 Effect on infarct volume 2 weeks post-stroke

To evaluate the effect of hybrid hydrogels on post-stroke brain repair, hybrid hydrogels (F1, F2, F3 and F6) were implanted into the infarct cavity 5 days after stroke. The infarct volume was assessed 2 weeks post-stroke using cresyl violet staining (**Fig. 2A**). Photothrombotic stroke produced a distinct focal lesion in the primary motor cortex of the left hemisphere (**Fig. 2B**). Intracerebral implantation of hydrogels resulted in differential effects on infarct volume. The optimised F6 hydrogel significantly reduced infarct size compared to no gel controls, indicating a measurable neuroprotective effect of the optimised formulation. No significant effect was seen in other hydrogel groups. This suggests that hydrogel composition critically influences neuroprotective outcomes.

**Figure 2:**
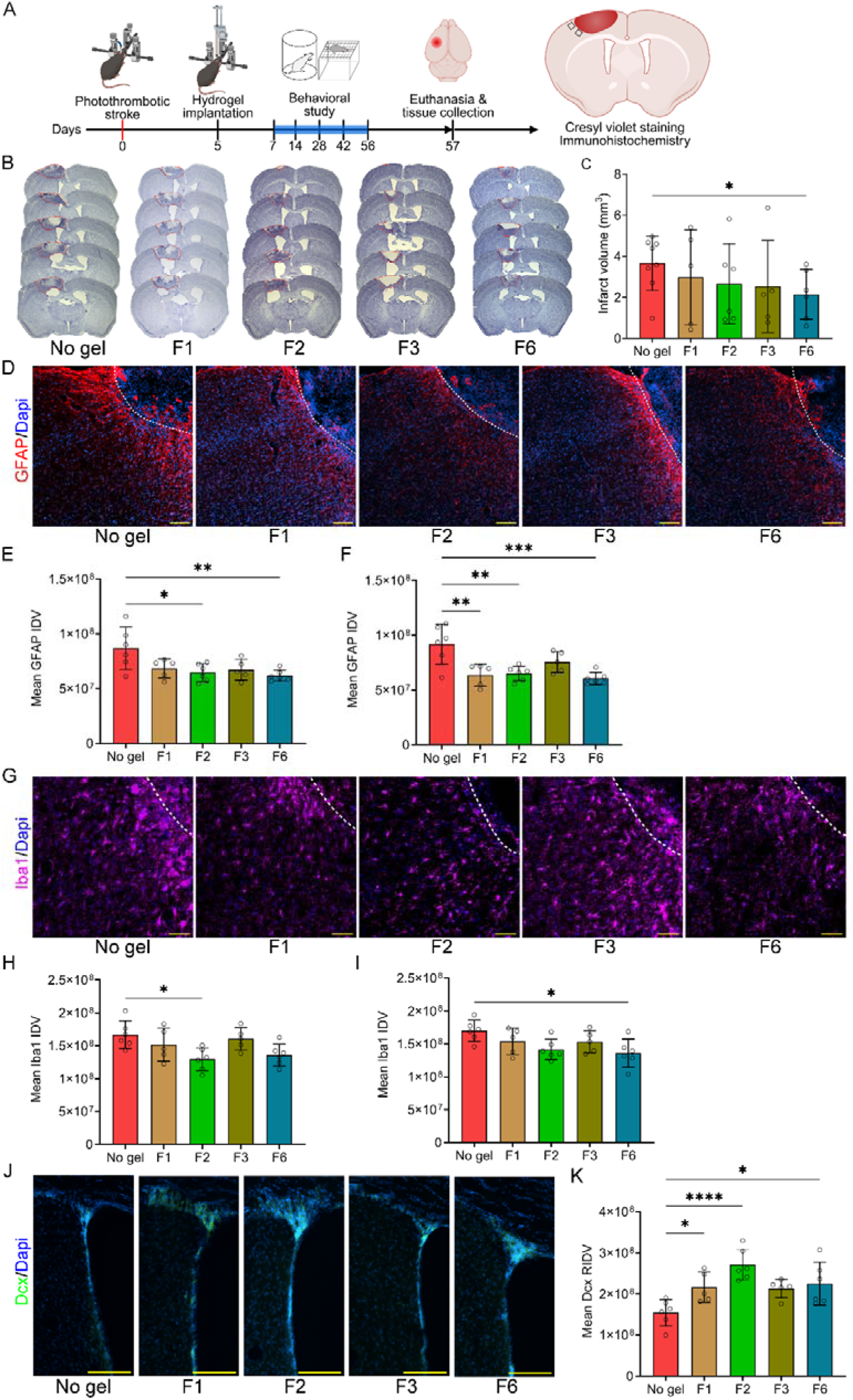
Hybrid hydrogels ameliorate infarct volume, reactive astrogliosis, microgliosis while promoting neurogenesis at 2 weeks post-stroke. (A) Experimental design. Mice underwent photothrombotic stroke and 5-days after stroke were given either one of these treatments: no gel (n=6), F1 (n=5), F2 (n=6), F3 (n=5) and F6 (n=6) through intracerebral injection directly into the infarct cavity. Mice were perfused 14-days after stroke, and brains collected for staining. (B) Representative Cresyl violet stained images from all treatment groups. (C) Quantification of infarct volumes at 14-days after stroke. (D) Representative fluorescent images of GFAP (+ve) astrocytes (shown in red) in the peri-infarct area at 2-weeks after stroke from all treatment groups. Quantification of mean integrated density values (IDV) of GFAP (+ve) astrocytes at (E) the cortical layer 2/3 peri-infarct, and (F) cortical layer 5 peri-infarct. (G) Representative fluorescent images of Iba1 (+ve) microglia (shown in purple) in the peri-infarct area from all treatment groups. Quantification of mean integrated density values (IDV) of Iba1 (+ve) microglial cells at (H) the cortical layer 2/3 peri-infarct, and (I) cortical layer 5 peri-infarct. (J) Representative fluorescent images of Dcx (+ve) neuroblasts (shown in green) at ipsilateral sub-ventricular zone (SVZ) from all treatment groups. (K) Quantification of raw integrated density values (RIDV) of Dcx (+ve) neuroblasts at ipsilateral SVZ. Data are expressed as mean ± SD and dots represent individual animals. Data were analysed by one-way ANOVA followed by Tukey’s post-hoc comparisons. Asterisks (*) represented significant values, * = P ≤ 0.05, ** = P ≤ 0.01 and *** = P ≤ 0.001.

### 3.3 Effect on reactive astrogliosis 2 weeks post-stroke

Stroke causes morphological changes in astrocytes including proliferation and hypertrophy, a process known as reactive astrogliosis [50]. Reactive astrogliosis contributes to glial scar formation in the peri-infarct region. The effect of hybrid hydrogels on reactive astrogliosis was assessed by quantifying the expression of GFAP in cortical layers 2/3 and 5 in the peri-infarct and lateral (distal) regions 2 weeks after stroke [51, 52]. Stroke induced a marked increase in GFAP expression in the peri-infarct region with a gradual decrease in expression in more lateral regions (**Fig. 2D**). Treatment with F6 hydrogel resulted in significant reduction of GFAP expression in both cortical layers 2/3 and 5 in the peri-infarct region (**Fig. 2E-F**), indicating attenuation of reactive astrogliosis in the peri-infarct region. Similar but less consistent effects were observed with F1 and F2, while F3 showed no significant impact. These findings suggest that specific hydrogel compositions can modulate astrocytic responses after injury. No significant difference was observed among the groups in the lateral cortex (**Fig. S3**).

### 3.4 Effect on microglial activation 2 weeks post-stroke

Similar to reactive astrocytes, stroke causes microglial activation in the peri-infarct region, promoting a proinflammatory response that worsens stroke pathology during the subacute and chronic phase [53–55]. To evaluate the effect of hybrid hydrogels on post-stroke microglial activation, the expression of Iba1 was evaluated in cortical layers 2/3 and 5 in the peri-infarct area 2 weeks after stroke [14]. Iba1 expression was significantly elevated in the peri-infarct region, reflecting microglial activation (**Fig. 2G**). Treatment with F2 and F6 hydrogels reduced Iba1 expression in a layer-specific manner, indicating partial suppression of microglial activation (**Fig. 2H-I**). No decrease in Iba1 expression was observed in the F1 or F3 hydrogel groups. These results suggest that hydrogel composition influences inflammatory responses in the peri-infarct region.

### 3.5 Effect on neurogenesis 2 weeks post-stroke

Stroke causes neuroblast proliferation in the subventricular zone (SVZ) and newly generated neuroblasts migrate towards the infarct area to promote regeneration [56]. The effect of hybrid hydrogels on post-stroke neurogenesis was assessed by quantifying the expression of Dcx in the ipsilateral SVZ (**Fig. 2J)**. Treatment with the F2 hydrogel led to a marked increase in Dcx expression compared to no gel controls, suggesting a potential role of SF in neuroblast proliferation (**Fig. 2K)**. The F1 and F6 hydrogels also significantly enhanced Dcx expression, indicating the capacity of these biomaterials to promote neuroblast proliferation after stroke.

### 3.6 Effect on behavioural function after stroke

Based on the effects on infarct volume, reactive astrogliosis and microgliosis, the F6 hydrogel was further assessed for post-stroke functional recovery. For this experiment, F6 hydrogel or F6 hydrogel combined with BDNF were injected into the infarct cavity 5 days after stroke and mice were tested for sensorimotor coordination using grid walk test and forelimb asymmetry using cylinder test for 8 weeks after stroke [15, 41]. For both tasks, a baseline measurement was performed 1 week prior to stroke surgery with behavioural assessments taken at 1-, 2-, 4-, 6– and 8-week post-stroke (**Fig. 3A**). Baseline assessment showed no difference among the treatment groups before the stroke. Animals exhibited a decreased contralateral limb use in cylinder test and an increased number of foot-faults in grid walk test from 1-week post-stroke until 8-week post-stroke, indicating persistent motor deficits (**Fig. 3B-C**). Treatment with F6 alone did not significantly improve functional outcomes. However, F6 combined with BDNF resulted in a significant improvement in both grid-walking and cylinder tests, with recovery emerging from 2–4 weeks and sustained over 8 weeks. These findings demonstrate that F6 hydrogel can act as an effective delivery platform for neurotrophic factors, translating biological effects into functional recovery

**Figure 3:**
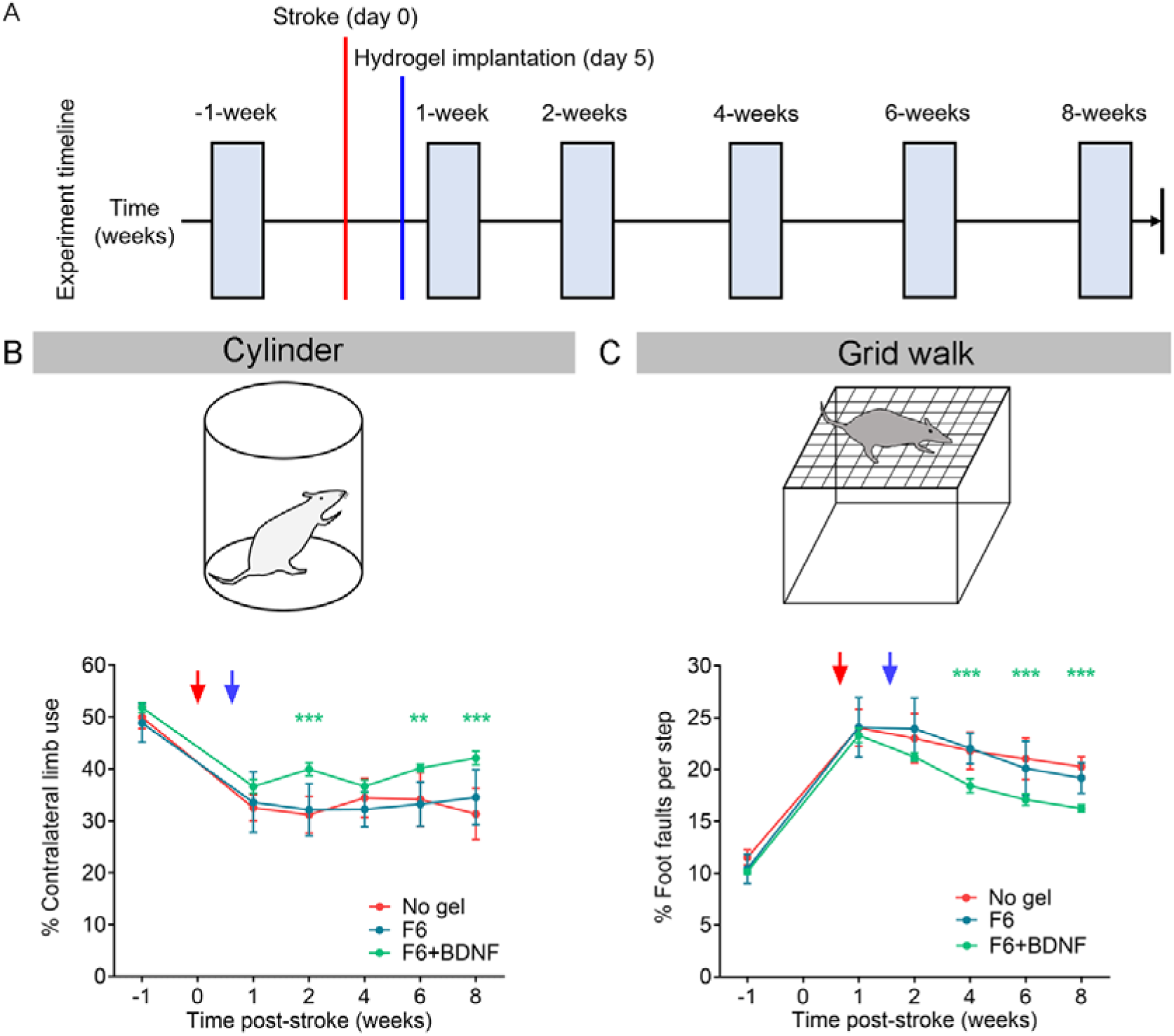
BDNF delivery from F6 hydrogel promotes behavioural recovery after stroke. (A) Experimental timeline for behavioural assessment. Mice were given one of the following treatments: no gel (n=8), F6-gel (n=9), F6+BDNF (n=9), through intracerebral injection directly into the infarct cavity. Mice were tested 1 week before stroke (baseline), and then subsequently at 1-, 2-, 4-, 6-, and 8-weeks after stroke to evaluate behavioural recovery on grid walk and cylinder tasks. (B) Evaluation of contralateral forelimb asymmetry using cylinder test, and (C) assessment of % foot faults per step for contralateral forelimb using grid walk test. Red and blue arrows represent the day of stroke surgery and the day of hydrogel implantation, respectively. Data are presented as mean ± SD. Data were analysed by two-way ANOVA followed by Tukey’s test for post-hoc comparisons. Asterisks (*) represented significant values, ** = P ≤ 0.01, *** = P ≤ 0.001.

### 3.7 Effect on infarct volume 8 weeks post-stroke

The effect of F6 hydrogel and F6+BDNF treatment on infarct size was assessed using cresyl violet staining 8 weeks post-stroke (**Fig. 4A**). Animals from all treatment groups showed visible lesions in cresyl violet staining (**Fig. 4B**). However, no significant difference in the infarct volume was observed among the treatment groups, suggesting that hydrogels had no effect on long-term infarct size (**Fig. 4C**).

**Figure 4:**
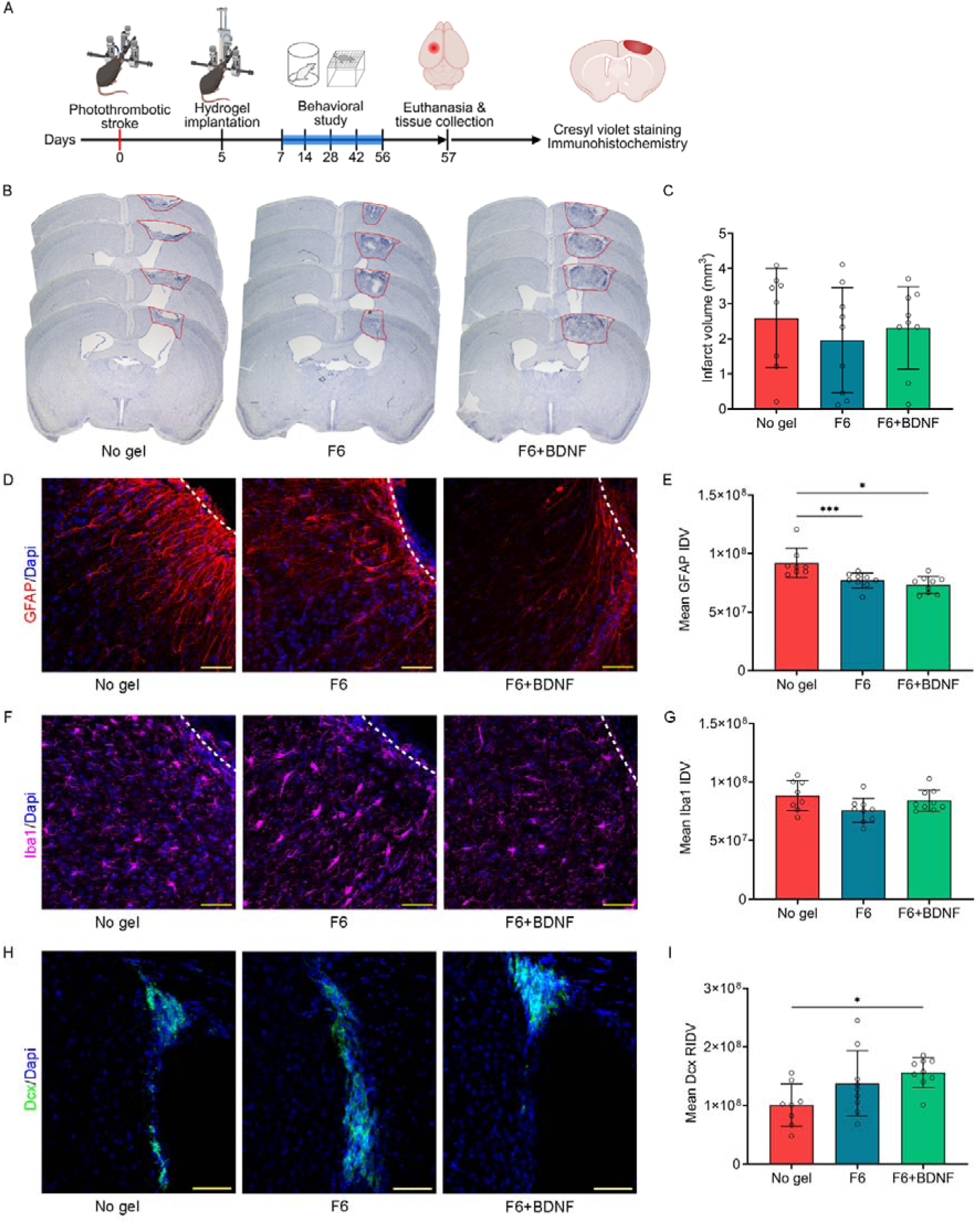
F6 hydrogel with BDNF attenuates reactive astrogliosis while promoting neurogenesis at 8 weeks post-stroke. (A) Experimental design. Mice underwent photothrombotic stroke and 5-days after stroke were given either one of these treatments: no gel (n=8), F6 (n=9), and F6+BDNF (n=9) through intracerebral injection directly into the infarct cavity. After completing 8-weeks behavioural data collection, mice were perfused, and brains collected for staining. (B) Representative Cresyl violet stained images from all treatment groups. (C) Quantification of infarct volumes 8 weeks after stroke. (D) Representative fluorescent images of GFAP (+ve) astrocytes (shown in red) in the peri-infarct area 8 weeks after stroke from all treatment groups. (E) Quantification of mean integrated density values (IDV) of GFAP (+ve) astrocytes in peri-infarct tissue. (F) Representative fluorescent images of Iba1 (+ve) microglia (shown in purple) in the peri-infarct area from all treatment groups. (G) Quantification of mean IDV of Iba1 (+ve) microglial cells in peri-infarct area. (H) Representative fluorescent images of Dcx (+ve) neuroblasts (shown in green) at ipsilateral sub-ventricular zone (SVZ) from all treatment groups. (I) Quantification of raw integrated density values (RIDV) of Dcx (+ve) neuroblasts at ipsilateral SVZ. Data are expressed as mean ± SD and dots represent individual animals. Data were analysed by one-way ANOVA followed by Tukey’s post-hoc comparisons. Asterisks (*) represented significant values, * = P ≤ 0.05, and *** = P ≤ 0.001.

### 3.8 Effect on reactive astrogliosis 8 weeks post-stroke

The effect of both F6 hydrogel and F6+BDNF on reactive astrogliosis was assessed 8 weeks post-stroke using GFAP staining. as neurotrophins can change the reactive properties of the peri-infarct tissue after stroke [16, 57]. All treatment groups demonstrated elevated expression of GFAP in the peri-infarct tissue, indicating a prolonged and persistent upregulation in reactive astrocytes 8 weeks post-stroke (**Fig. 4D**). Treatment with both F6 hydrogel and F6+BDNF significantly decreased GFAP expression in peri-infarct tissue (**Fig. 4E**), suggesting dampening of reactive astrogliosis in the chronic stage after stroke.

### 3.9 Effect on microglial activation 8 weeks post-stroke

The effect of both F6 hydrogel and F6+BDNF on activation of microglial cells was assessed by quantifying Iba1 expression in the peri-infarct tissue 8 weeks after stroke (**Fig. 4F**). No difference in Iba1 expression was observed in the peri-infarct area among different intervention groups, indicating that early changes in microglial reactivity have been normalised by 8 weeks post-stroke (**Fig. 4G**).

### 3.10 Effect on neurogenesis 8 weeks post-stroke

To assess the long-term effect of F6 hydrogel and F6+BDNF on neurogenesis, neuroblasts were quantified using Dcx staining 8 weeks following stroke (**Fig. 4H**). Treatment with F6 showed no significant effect on Dcx expression, however F6+BDNF treatment demonstrated a significant increase in Dcx expression in the SVZ (**Fig. 4I**), suggesting that BDNF promoted long term neurogenesis after stroke.

## 4 Discussion

In the present study, we developed a thermoresponsive hybrid hydrogel system composed of CS, β-GP, SF, PVA, and PVP, and evaluated its potential to modulate the post-stroke microenvironment and promote functional recovery. By combining natural and synthetic polymers, we demonstrated that polymer composition can be systematically tuned to control injectability, gelation kinetics, mechanical properties, and biological functionality, key parameters for biomaterials intended for brain repair [58, 59].

CS/β-GP hydrogels are well known for their thermoresponsive gelation, driven by a combination of electrostatic interactions, hydrogen bonding and hydrophobic interactions [44]. In this study, incorporation of additional polymers significantly modulated the gelation kinetics and mechanical properties. SF increased gelation temperature and reduced stiffness, whereas PVA and PVP promoted faster gelation and enhanced viscoelastic properties. These findings highlight that relatively small changes in polymer composition can substantially alter intermolecular interactions within the hydrogel network, resulting in tuneable physicochemical behaviour. Importantly, the optimised formulation (F6) demonstrated gelation near physiological temperature while maintaining suitable mechanical properties for brain tissue, a critical consideration given that neural tissue is highly sensitive to mechanical mismatch, with substrate stiffness known to directly regulate astrocyte activation, neural differentiation, and stem cell fate [20, 60, 61].

Hydrogel microarchitecture is a key determinant of cellular infiltration and molecular transport [62]. In the present study, all formulations showed interconnected microporous structures, which are known to facilitate cell migration and diffusion of bioactive molecules [62]. Previous studies have demonstrated that microporous hydrogels can reduce glial scar formation and enhance tissue remodelling after stroke by enabling cell infiltration and modulating local signalling gradient [14, 15]. Consistent with these reports, the hydrogels developed in this study were associated with reduced reactive astrogliosis in the peri-infarct region. Mechanistically, this effect is likely mediated by a combination of biomechanical cues (e.g., matrix and porosity), altered cell-matrix interactions, and modulation of ECM-like properties as well as attenuation of inflammatory signalling [14, 15].

Reactive astrocytes are known to produce chondroitin sulphate proteoglycans, ephrin-A5, and other inhibitory extracellular matrix components in the peri-infarct region that contribute to glial scar formation, inhibit axonal regeneration and neural plasticity [19, 63–65]. Therefore, pharmacological agents that decrease peri-infarct reactive astrogliosis are being trialled preclinically and may create a more permissive environment for neural repair [51, 52, 66].

Biodegradability is essential for biomaterial translation, particularly in the brain where long-term foreign body presence can exacerbate inflammation [67, 68]. In the current study, all hybrid hydrogels demonstrated progressive enzymatic degradation, consistent with the known lysozyme-mediated breakdown of chitosan into biocompatible metabolites [69]. The degradation profile suggests that the materials can provide temporary structural support during the critical window of post-stroke repair while gradually being resorbed, thereby minimizing chronic foreign body responses. Similar degradation behaviour has been reported for CS/β-GP hydrogels in vivo [70].

*In vitro* cytotoxicity assays confirmed that all formulations were well tolerated by PC12 cells, indicating cytocompatibility [71–73]. Notably, some formulations (F3 and F6) promoted increased cell viability, suggesting that polymer composition may actively influence cellular responses, potentially through modulation matrix stiffness, surface properties or protein adsorption [68]. However, the mechanisms underlying these effects require further investigation.

During the *in vivo* study, the optimised hydrogel (F6) reduced infarct volume at 2 weeks post-stroke and significantly attenuated reactive astrogliosis. In addition, F6 hydrogel decreased microglial activation indicating modulation of pro-inflammatory signalling during the subacute phase of stroke [53–55, 74]. Together, these findings suggest that the F6 hydrogel modulates key cellular components of the post-stroke microenvironment, including astrocytes and microglia, thereby shifting the injury environment from a proinflammatory, inhibitory state towards a more permissive and pro-regenerative niche [75, 76].

In the present study, the observed increase in neurogenesis in the SVZ further supports the pro-regenerative potential of the F6 hydrogel. Endogenous neural progenitor cells are known to proliferate and migrate toward injury sites after stroke, although their survival and integration are limited [56].

Biomaterial scaffolds can enhance this process by providing structural and biochemical support [15, 17, 77]. While this effect may be partially attributed to the presence of bioactive motifs within SF, such as RGD sequences [78], it is also likely influenced by the overall matrix environment, including mechanical and structural cues that support neural progenitor cell proliferation and migration [60, 79].

Incorporation of BDNF within F6 hydrogel significantly enhanced functional recovery over 8 weeks. The role of BDNF is well established for enhancing neural plasticity and functional recovery after stroke [16, 80], but its clinical application is limited by rapid degradation and poor retention in tissue [80, 81]. Hydrogel-based delivery systems can improve the stability, retention and spatial localization of BDNF, enabling sustained activation of pro-plasticity signalling pathways and improved functional outcomes [16, 58, 82]. Here, the F6 hydrogel likely acts as a local reservoir, enhancing retention and sustained availability of BDNF within the peri-infarct tissue. However, the release kinetics and dose-response relationship of BDNF from the F6 hydrogel were not assessed in this study, limiting mechanistic interpretation and representing an important area for future investigation.

Despite these promising findings, several limitations should be acknowledged. First, the molecular mechanisms underlying hydrogel-mediated modulation of astrocytes and microglia were not directly investigated in this study. Second, quantitative characterisation of hydrogel mechanical properties relative to native brain tissue was not performed, which is important for establishing the structure – function relationship and optimising material design [60]. Third, BDNF release profiles and dose–response relationships were not evaluated, limiting mechanistic interpretation of the functional recovery results. Finally, long-term in vivo degradation of the biomaterial and related immune responses require further assessment to support translation to a therapeutic setting [67, 68].

Importantly, comparison with established biomaterial systems highlights both the potential and limitations of the present approach. Hyaluronic acid-based hydrogels and microporous annealed particle (MAP) systems have demonstrated efficacy in promoting tissue repair following stroke, in part due to their ability to mimic native ECM components [15, 83, 84]. While the hybrid thermoresponsive hydrogel described here offers advantages in terms of injectability and tunability, incorporation of defined ECM-mimetic or glycomimetic components may further enhance its biological performance.

Future studies should focus on elucidating the cellular and molecular mechanisms of hydrogel–tissue interactions, optimising mechanical and biochemical properties to better match brain tissue, and characterising controlled release of therapeutic factors. Comparative studies with established biomaterials, such as hyaluronic acid-based hydrogels, which have shown efficacy in stroke repair models [14, 15], would also strengthen the translational positioning of this system.

## 5 Conclusion

Biopolymer hydrogels, with or without additional bioactive molecules, have been studied extensively as a therapeutic strategy for brain repair following stroke. In the present study, we have examined biocompatibility and therapeutic effect of CS, SF, PVA, PVP and β-GP containing hybrid hydrogels using a photothrombotic model of ischemic stroke. Among the tested hydrogels, the optimised F6 hydrogel demonstrated promising bio-physicochemical including biocompatibility with brain tissue, rapid gelation, microporous architecture, and controlled biodegradation. Mechanistically, the hybrid hydrogel F6 modulated reactive astrogliosis, reduced microglial activation and enhanced neurogenesis after stroke. Additionally, combining neurotrophic factor, BDNF with F6 hydrogel promoted functional recovery after stroke. Further studies are recommended to characterise the potential of this hydrogel for sustained release of therapeutics and other growth factors.

## 8 Authors contribution

MHB: conceptualisation, methodology, investigation, analysis, writing-original first draft & editing; EKG: investigation, analysis; UZM: investigation, analysis; SFRH: writing-review & editing; MAA: conceptualisation, supervision, visualisation, writing-review & editing; ANC: conceptualisation, supervision, visualisation, writing-review & editing.

