## Supplementary Materials for "Thermoresponsive chitosan/silk fibroin/PVA/PVP hydrogel loaded with BDNF promotes functional recovery after stroke"

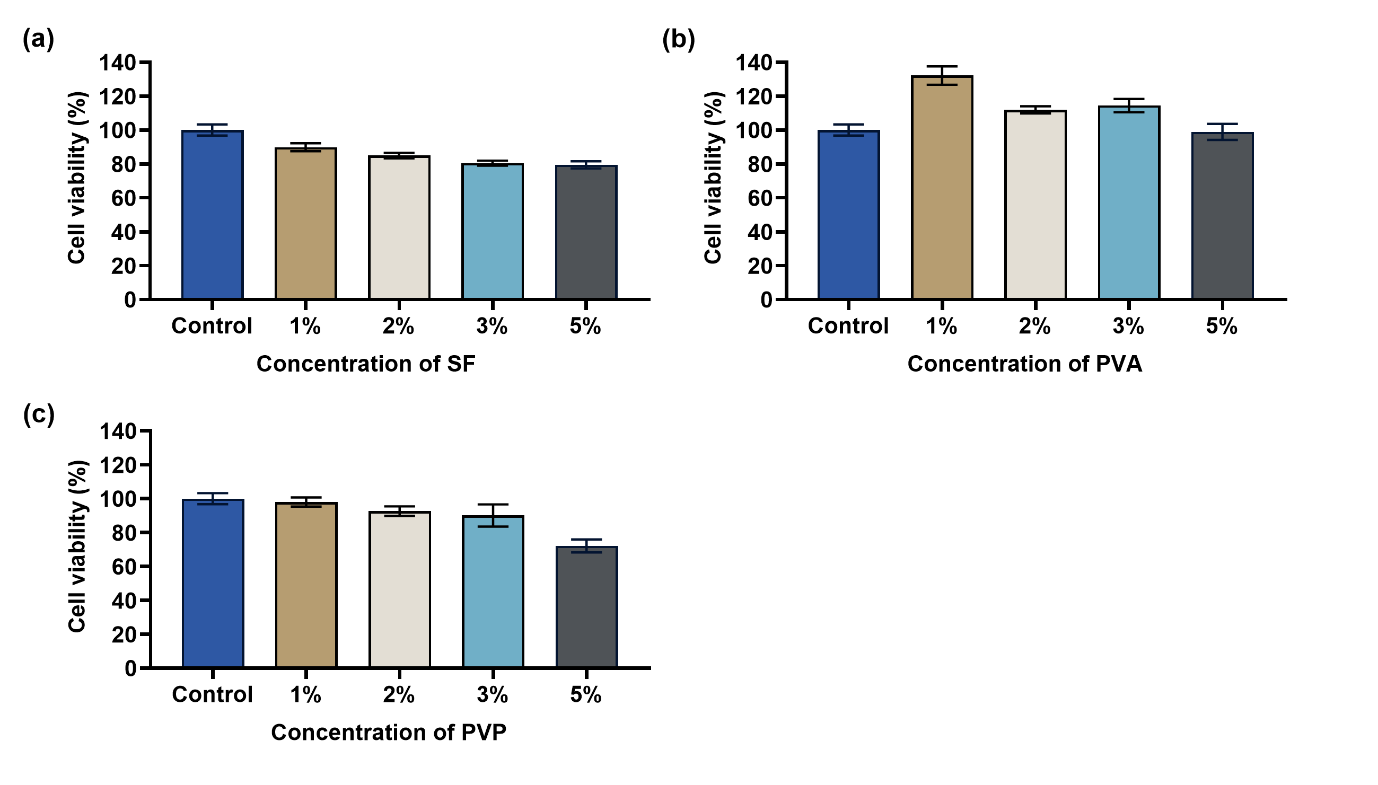

Figure S1: Cell viability of PC12 cells cultured with silk fibroin (SF), polyvinyl alcohol (PVA) and polyvinyl pyrrolidone (PVP) at 24 h measured by MTT assay.

(a) 1, 2, 3, and 5% w/v SF solution, (b) 1, 2, 3, and 5% w/v PVA solution, and (c) 1, 2, 3, and 5% w/v PVP solution. Data are expressed as mean ± SD for n=3.

### Fourier transform infrared (FTIR) characterization of the hybrid hydrogels

**Method:** The FTIR spectroscopy (ALPHA FTIR, Bruker, USA) was used to characterise the chemical bonds present in the hybrid hydrogels. The spectrums were obtained in ATR mode over the wavelength range between 400 – 4000 cm^-1^ with a spectral resolution of 4 cm^-1^.

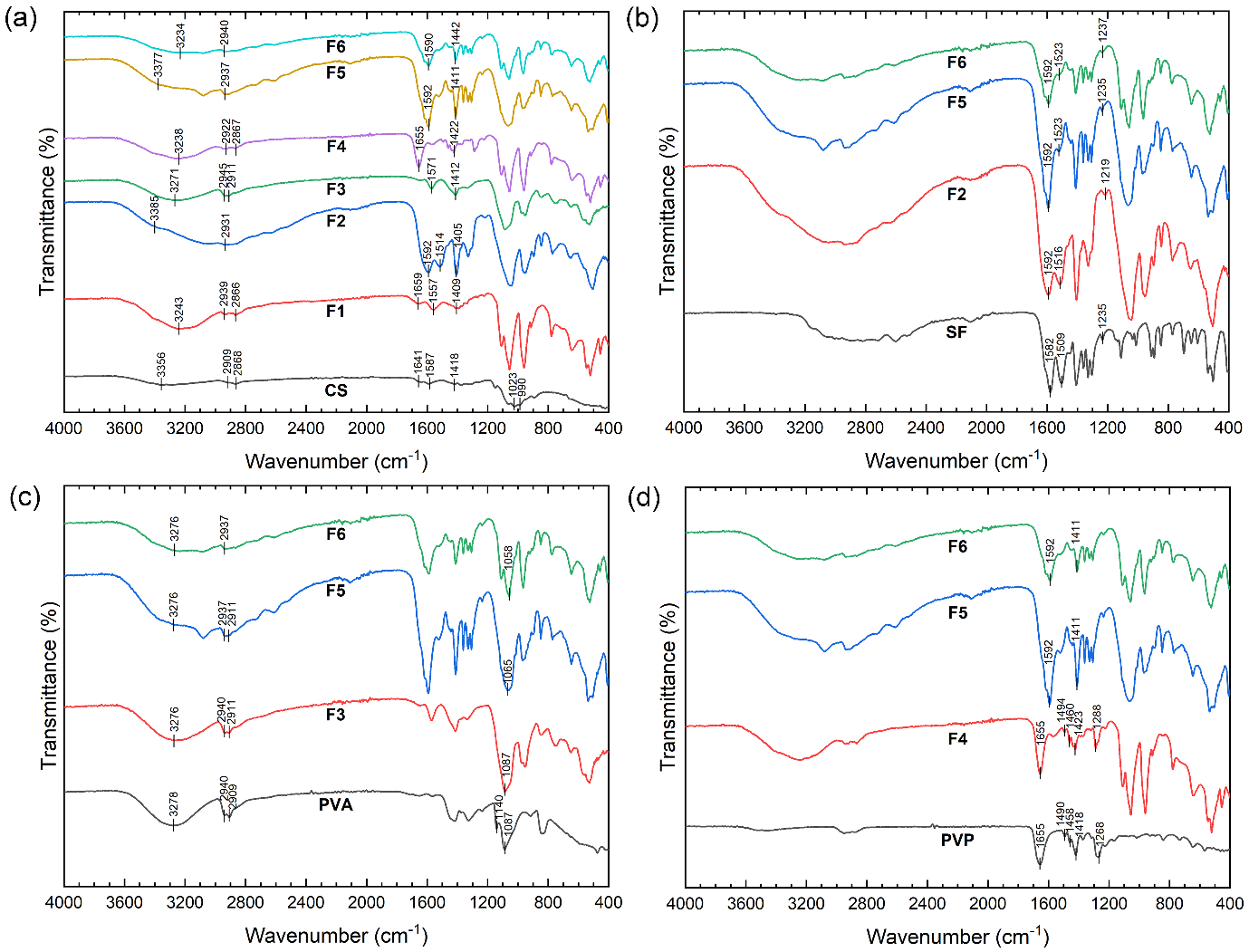

Figure S2: FTIR spectra of hybrid hydrogels along with (a) chitosan, (b) silk fibroin, (c) polyvinyl alcohol and (d) polyvinyl pyrrolidone. CS: chitosan, SF: silk fibroin, PVA: polyvinyl acetate, PVP: polyvinyl pyrrolidone, F1: 0.75% chitosan / 3% β-GP, F2: 0.75% chitosan / 3% β-GP/ 3% SF, F3: 0.75% chitosan / 3% β-GP/ 2% PVA, F4: 0.75% chitosan / 3% β-GP/ 1% PVP, F5: 0.75% chitosan / 3% β-GP/ 3% SF/ 2% PVA/ 1% PVP, F6: 0.75% chitosan / 3% β-GP/ 2% SF/ 1% PVA/ 0.5% PVP.

**Results:** The chemical interactions among CS, SF, PVA, PVP and β-GP were investigated using FTIR spectroscopy (**Fig. S2)**. The FTIR spectrum of pure CS showed a broad shoulder between 3400-3200 cm^-1^, corresponding to O-H stretching overlapped with the N-H stretching. Peaks at 2909 and 2868 cm^-1^ indicating C-H stretching, while peaks at 1641 cm^-1^ (C=O bending), 1587 cm^-1^ (N-H bending), and 1418 cm^-1^ (C-CH_3_ symmetric deformation) were also observed, in agreement with previous reports [48]. Pure SF displayed characteristic peaks at 1582 and 1509 cm^-1^, corresponding to C=O stretching and N-H bending associated with β-sheet structures. An additional peak at 1235 cm^-1^ was attributed to C-N and N-H vibrations from random coil and α-helix structures [27]. The FTIR spectrum of pure PVA showed a broad peak between 3450-3200 cm^-1^, corresponding to O-H stretching, peaks at 2940 and 2908 cm^-1^ for C-H stretching from alkyl groups, and a prominent peak at 1140 cm^-1^for C-O stretching. For pure PVP, characteristic peaks included a strong C=O stretch (amide I) at 1655 cm^-1^ , aromatic to C-H deformation at 1490 and 1458 cm^-1^, C-N stretching in the pyrrolidone structure at 1288 cm^-1^ [49].

In the hybrid hydrogels, characteristic peaks of CS were retained, with a shift in the O-H stretching band in F2 due to SF incorporation. Minor shifts at 1647, 1587, and 1418 cm⁻¹ indicated interactions with other polymers (**Fig. S2a**). SF-specific peaks were observed in F2, F5, and F6 hydrogels, with the C=O peak shifting from 1582 to 1592 cm⁻¹ and N–H bending shifting from 1509 to 1516 cm⁻¹ (F2) and 1523 cm⁻¹ (F5 and F6; **Fig. S2b**). Similarly, PVA peaks were identifiable in F3, F5 (**Fig. S2c**), and F6, and PVP peaks were evident in F4, F5, and F6 with minor shifts indicating polymer–polymer interactions (**Fig. S2d**).

Table S1: Animal sample size (n) for all experiments

| Study length | Treatment group | Stroke | Total enrolled | Death | Exclusion* | Final sample size |
| --- | --- | --- | --- | --- | --- | --- |
| 2-weeks | No gel | + | 6 |  |  | 6 |
|  | F1 | + | 6 |  | 1 | 5 |
|  | F2 | + | 6 |  |  | 6 |
|  | F3 | + | 6 | 1 |  | 5 |
|  | F6 | + | 6 |  |  | 6 |
| 8-weeks | No gel | + | 9 |  | 1 | 8 |
|  | F6 | + | 9 |  |  | 9 |
|  | F6 + BDNF | + | 9 |  |  | 9 |

*Animals with an infarct volume smaller than 0.2 mm^3^ were excluded from data analysis.

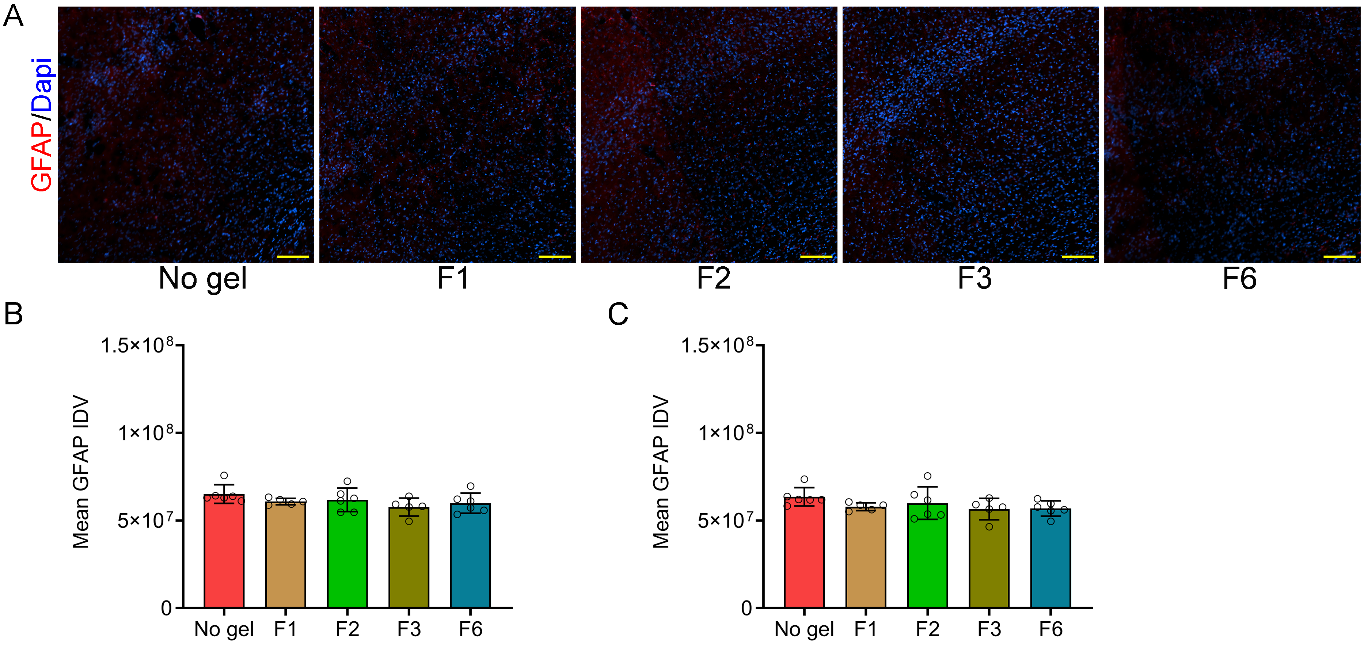

Figure S3: GFAP expression in the lateral cortical regions were similar across treatment groups at 2-weeks post-stroke.

(a) Representative fluorescent images of GFAP (+ve) astrocytes (shown in red) in the lateral area at 2-weeks after stroke from all treatment groups: no gel (n=6), F1 (n=5), F2 (n=6), F3 (n=5) and F6 (n=6). Scale bar = 50 µm. All sections have been counter stained with DAPI (shown in blue). Quantification of mean integrated density values (IDV) of GFAP (+ve) astrocytes at (b) the cortical layer 2/3, and (c) cortical layer 5. Data are expressed as mean ± SD and dots represent individual animals. Data were analysed by one-way ANOVA with Tukey’s test for post hoc comparisons.
